# GenNA: Conditional generation of nucleotide sequences guided by natural-language annotations

**DOI:** 10.64898/2026.04.22.720063

**Authors:** Yi Shen, Guangshuo Cao, Jianghong Wu, Dijun Chen, Cong Feng, Ming Chen

**Author notes:** Correspondence (Cong Feng), (Ming Chen). These authors contributed equally to this work (Yi Shen, Guangshuo Cao).

## Abstract

Deciphering the mapping between linear biomolecular sequences and complex biological functions remains a central challenge in genomics. Although existing generative nucleotide language models have made substantial progress in modeling sequence distributions, they generally lack explicit access to high-level biological semantics, limiting their ability to support semantics-guided conditional generation. To address this limitation, we present GenNA, a generative nucleotide foundation model guided by natural-language annotations. GenNA is pretrained on a multimodal nucleotide-text corpus spanning 2,221 eukaryotic species and comprising approximately 416 billion characters, and learns the relationships between sequence patterns and functional annotations within a unified autoregressive framework. Systematic evaluations show that, even without explicit supervision from biological rules, GenNA yields distinguishable perplexity scores in response to semantic mismatches between sequences and functional annotations, to different mutation types, and to perturbations of species labels. Moreover, across a range of natural-language-guided nucleotide generation tasks, the model produces sequences consistent with both target semantics and species context. Overall, GenNA provides a unified framework for natural-language-guided nucleotide modeling and conditional generation, and offers a feasible route toward integrating high-level functional descriptions with low-level sequence design.

## Main

Understanding how nucleotide sequences encode complex biological functions has long been a central question in genomics and synthetic biology^1^. In recent years, the rapid development of foundation models for protein and nucleic acid sequence modeling has transformed the view that biological sequences can be treated as a learnable language from a conceptual analogy into an operational computational paradigm^2^. In the protein domain in particular, large-scale self-supervised learning has shown that models can acquire representations associated with structure, fitness, and function purely from sequence statistics^3,4^. This progress has, in turn, directly motivated the extension of language-modeling approaches to DNA and RNA sequences.

Nucleotide foundation models have advanced along several related but distinct trajectories. For nucleotide sequence representation and prediction, encoder-based models such as DNABERT^5^ and Nucleotide Transformer^6^ have been followed by longer-context and sequence-to-function frameworks including DNABERT-2^7^, GENA-LM^8^, AlphaGenome^9^ and Nucleotide Transformer v3^10^, collectively showing that large-scale pretraining can capture regulatory, mutational and cross-species signals across increasingly long genomic contexts. For sequence generation, autoregressive models such as GenSLMs^11^, DNAGPT^12^ and HyenaDNA^13^, followed by later genome-scale models including megaDNA^14^, Evo^15^, Evo 2^16^, GENErator^17^ and ATGC-Gen^18^, have demonstrated that nucleotide sequences can be modeled and designed directly within generative frameworks.

However, the explicit use of natural-language annotations as the primary conditioning signal for nucleotide sequence generation remains comparatively underexplored. Previous approaches rely on structured labels or predefined attributes, whose limited expressivity struggles to capture complex, context-dependent biological functions. In contrast, natural language provides a vastly more flexible representation, more suitable for expressing human intent. In the absence of effective mechanisms to integrate such rich and heterogeneous information, most generative models rely mainly on sequence prefixes, local context, or specific structured attributes for conditional modeling. Even OmniNA^19^, despite jointly modeling nucleotide sequences and textual annotations, mainly emphasizes sequence-to-annotation generation and sequence understanding rather than annotation-conditioned sequence design.

Against this backdrop, we present GenNA, a generative nucleotide foundation model guided by natural-language annotations. GenNA adopts a decoder-only Transformer architecture^20,21^ configured with reference to Qwen3^22^, using a custom cross-modal BPE vocabulary^23^, and is pretrained on a unified nucleotide-text-structure-label corpus spanning 2,221 eukaryotic species and approximately 416 billion characters. Unlike existing models that are centered primarily on a single sequence modality or only weakly structured conditions, GenNA is explicitly designed to incorporate natural-language functional annotations, species context, and gene-related metadata into the autoregressive modeling process, enabling biological semantics to directly inform nucleotide sequence modeling and generation.

We further show that this multimodal training strategy enables the model to learn nucleotide sequence patterns while capturing the relationships among sequence features, functional semantics and species context. Without introducing explicit supervision from biological rules, GenNA exhibits higher perplexity to sequence-function mismatches, different mutation types, and perturbations of species labels. In addition, it supports controllable nucleotide generation under natural-language guidance, ranging from general open-ended scenarios to highly constrained tasks. Overall, GenNA fills a critical gap between sequence modeling and semantics-driven design in current nucleotide foundation models, providing a unified framework for natural-language-guided nucleotide understanding and generation.

## Results

### Architecture and pretraining of the GenNA foundation model

GenNA is a decoder-only generative foundation model for nucleotide sequences. Its network configuration follows the Qwen3 Transformer^22^ design paradigm, and the model was pretrained from scratch using a custom cross-modal vocabulary, resulting in approximately 3.6 billion parameters (Fig. 1a). Unlike nucleotide models that are primarily based on masked language modeling^24^, GenNA adopts an autoregressive framework to enable direct sequence generation. Building on this framework, GenNA incorporates natural-language prompts at the beginning of the sequence to provide biological information. This design extends the conditioning space of autoregressive generation by incorporating high-level biological semantics, including species identity, gene names and functional annotations, alongside local sequence context.

**Fig. 1:**
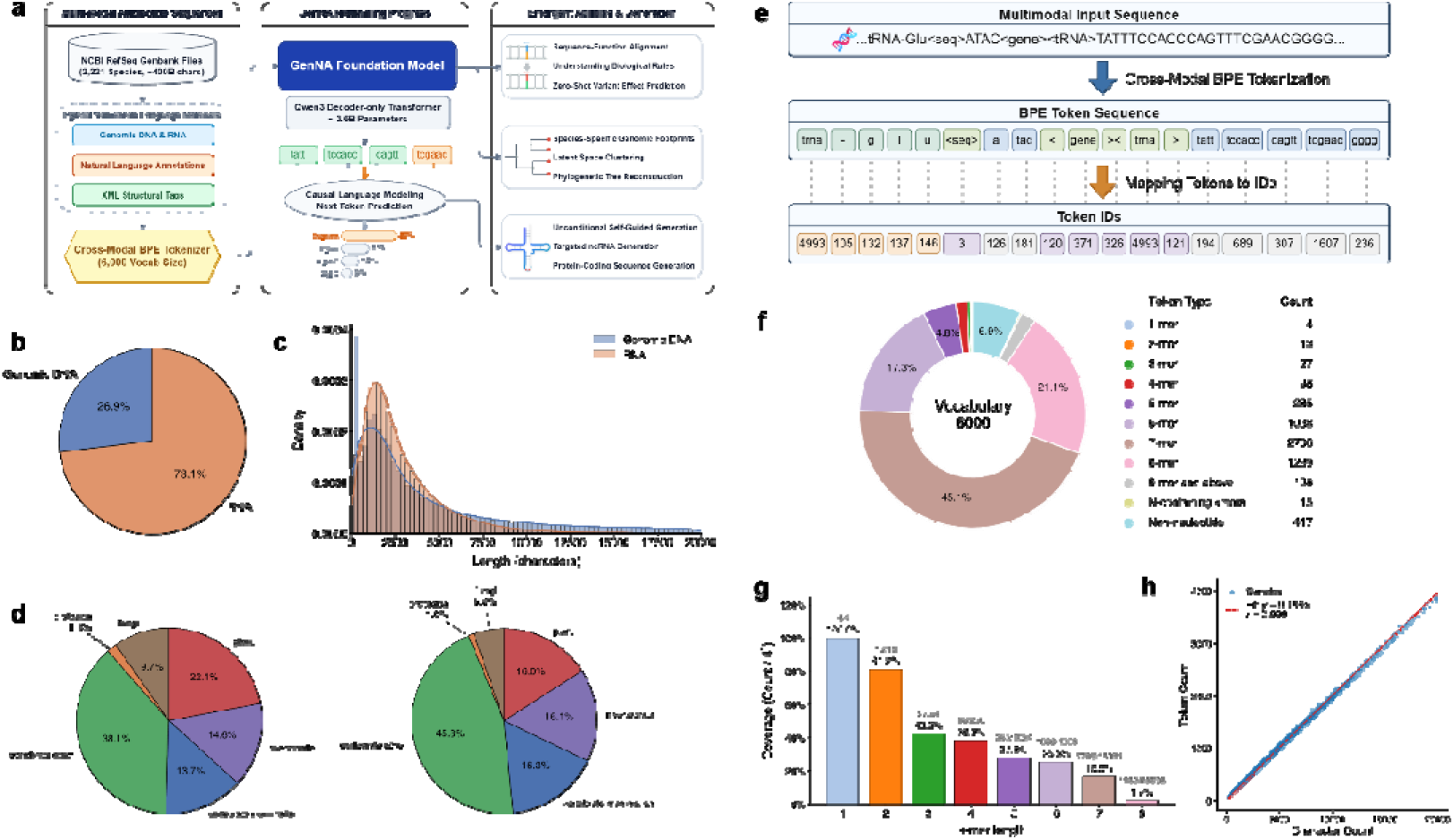
Architecture design and multimodal pretraining corpus of the GenNA foundation model. **a**, Overview of the GenNA pretraining pipeline, illustrating the integration of multimodal nucleotide sequences and natural-language annotations via a decoder-only Transformer. **b**, Proportion of genomic DNA and RNA in the pretraining corpus. **c**, Density distribution of sequence character lengths for DNA and RNA. **d**, Taxonomic distribution of the pretraining corpus across major eukaryotic lineages, measured by sequence count (left) and total character count (right). **e**, Schematic of the cross-modal BPE tokenization process mapping multimodal sequence and text inputs to unified token IDs. **f**, Composition of the 6,000-token BPE vocabulary categorized by token type and k-mer length. **g**, Sequence coverage rates achieved by different k-mer lengths within the vocabulary. h, Correlation between character count and token count, demonstrating sequence compression efficiency. The red dashed line represents the linear fit (Pearson’s).

The pretraining data were derived from the NCBI RefSeq database^25^. After curation, we constructed a multimodal corpus covering 2,221 eukaryotic species and totaling approximately 416 billion characters (Fig. 1b-d and Supplementary Table 1). In addition to nucleotide sequences from gene-associated regions, the corpus incorporates natural-language functional metadata and XML-style structured annotations (Supplementary Table 2). To handle this heterogeneous data type, we further trained a unified cross-modal BPE tokenizer^23^, allowing frequent functional phrases, tag fragments, and local nucleotide patterns to be mapped into a shared discrete token space (Fig. 1e-h). This facilitates the joint modeling of sequence patterns, functional semantics and structural boundary information within a unified representation space.

During training, we used a causal language modeling objective^20^, in which the model predicts the next token conditioned on the preceding context. GenNA was pretrained from scratch on a computing cluster composed of multiple NVIDIA A800 GPUs, with a maximum context length of 4,096 tokens, corresponding to roughly 20,000 raw characters. This context window is sufficient to cover the full context of a substantial fraction of real genes together with their associated annotations.

For reproducibility, we relased the model weights, training code, and inference code, together with Python scripts for processing full-format GenBank files and constructing aligned training corpora from standard GenBank records^26^. In addition, we provide a smaller 0.36B parameters version to facilitate efficient experimentation and development in resource-constrained settings.

### GenNA internalizes sequence-function relationships

The pretraining corpus contains a large number of paired nucleotide sequences and natural-language functional annotations, suggesting that the model may capture relationships between sequence patterns and functional semantics. To evaluate whether GenNA is genuinely sensitive to the consistency between sequences and their functional annotations, we first assessed its response to sequence-function mismatches using perplexity (PPL) and then examined attention scores in intermediate Transformer layers to assess how natural-language prompts contribute to autoregressive nucleotide generation.

While keeping the underlying nucleotide sequence completely unchanged, we systematically replaced the functional annotation in the input prompt. The results showed that, when the natural-language functional annotation was mismatched with the corresponding sequence, the model’s average PPL increased substantially (Fig. 2a,b). In addition, gene-annotation swapping across 15 genes from four families produced a structured ΔPPL pattern, suggesting that the model captures family-level functional relatedness (Supplementary Fig. 4).This elevated uncertainty in response to semantically inconsistent inputs indicates that GenNA does not rely solely on local nucleotide context but instead jointly incorporates functional-semantic information provided by the prompt.

**Fig. 2:**
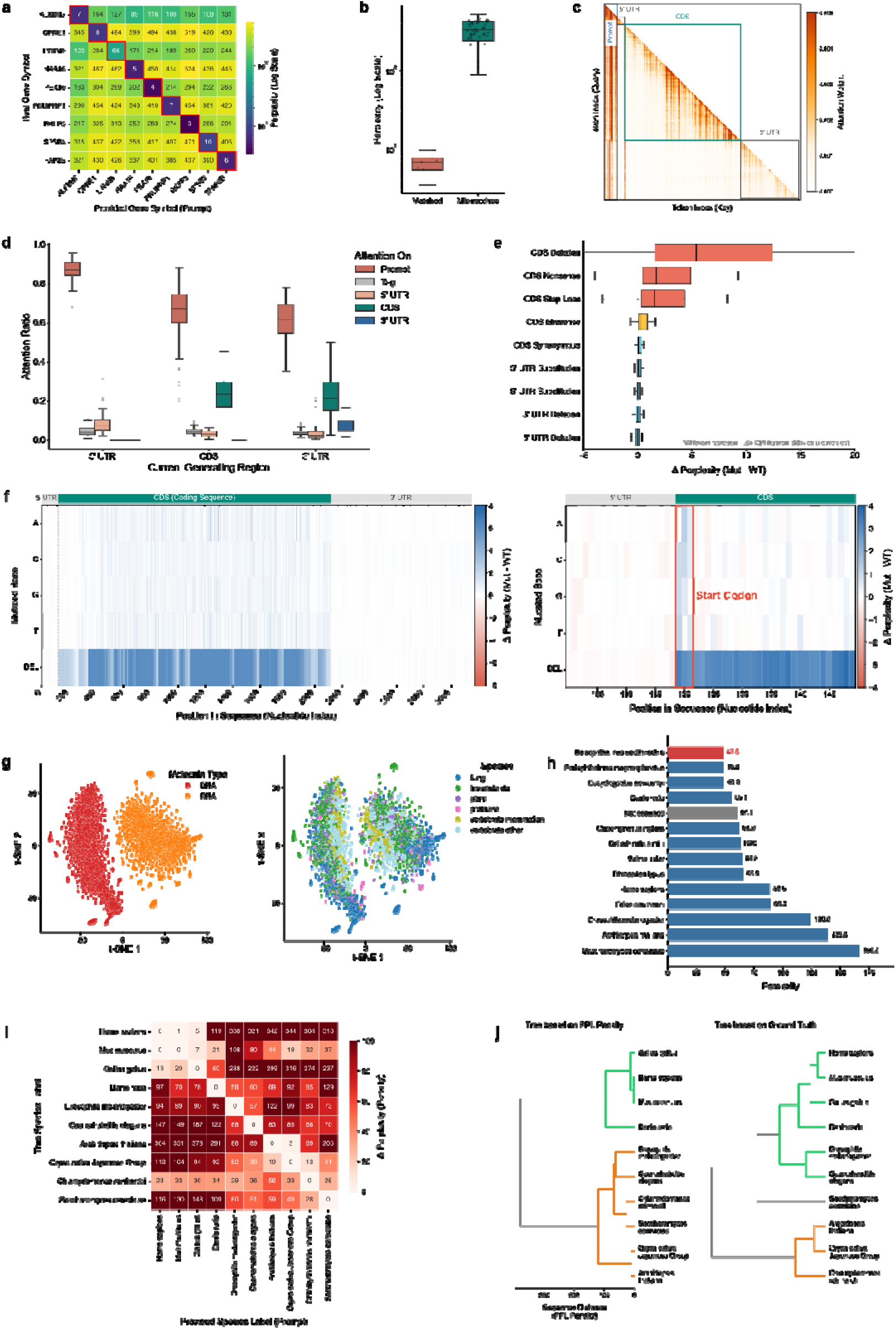
GenNA internalizes sequence-function mappings and evolutionary constraints. **a**, Perplexity (PPL) heatmap evaluating the alignment between real nucleotide sequences and provided gene anntotation prompts (genes). Warmer colors indicate higher ΔPPL. **b**, Perplexity for matched versus mismatched sequence-function pairs. **c**, Token-to-token self-attention map extracted from Transformer Layer 18 during autoregressive sequence generation. **d**, Dynamic attention ratio allocated to prompt tags versus upstream sequence components during the generation of the 5′ UTR, CDS, and 3′ UTR (*n* = 100 sequences). **e**, Zero-shot in silico variant effect prediction across different mutation types in protein-coding genes (*n* = 100 sequences). ΔPPL represents the difference in perplexity between mutant and wild-type sequences. **f**, High-resolution ΔPPL landscape along a representative coding sequence. **g**, t-SNE visualization of intermediate-layer sequence embeddings, colored by taxonomic group (left) and molecule type (right). **h**, Baseline perplexity for native sequences across 10 representative species. **i**, ΔPPL matrix induced by species prompt swapping. **j**, Topological comparison between the hierarchical clustering tree derived from GenNA’s pairwise ΔPPL (left) and the biological reference phylogenetic tree (right).

To further investigate this phenomenon at the level of internal representations, we analyzed the token level attention scores in Transformer layer 18^27^. Using the *Spodoptera litura* glutaminase liver isoform, mitochondrial isoform X1 for example, we observed that even when the generation position was distant from the prompt region, the model continued to attend to functional annotations and structural labels throughout the generation process (Fig. 2c). Across 100 randomly sampled sequences, we observed that during coding sequence (CDS) generation, prompt tokens accounted for an average of 64.4% of the attention weight, whereas relatively less attention was assigned to the upstream 5’ UTR (Fig. 2d). This pattern is consistent with the interpretation that prompt tokens continue to provide conditioning constraints throughout the generation process. Prompt ablation further supported the interpretation that functional annotation provides the dominant semantic signal (Supplementary Fig. 3).

Taken together, these results indicate that GenNA captures relationships between natural-language functional annotations and nucleotide sequences. Functional prompts are therefore not merely superficial auxiliary information in the input, but actively participate in both the probabilistic evaluation and the generation of the entire conditioned sample.

### Zero-shot prediction of mutation effects

To evaluate the sensitivity of GenNA to single-nucleotide perturbations in functional sequences, we performed in silico mutational scanning on protein-coding genes and compared the perplexity changes induced by different mutation types (Fig. 2e,f). Within the evaluation framework used in this study, ΔPPL can be interpreted as the model’s response how far a mutated sequence deviates from the natural distribution^28,29^. It therefore provides a useful proxy for comparing the relative impact of different classes of perturbations on the model’s internal assessment of sequence function.

The results revealed a clearly stratified pattern across mutation categories. In general, single-nucleotide mutations occurring in non-coding regions (such as 3′ and 5′ UTR substitutions or deletions), as well as synonymous substitutions within the coding sequence (CDS), caused only modest changes in ΔPPL (median values ranging from -0.015 to 0.076). This suggests that the model is relatively tolerant of local perturbations that do not alter the encoded product (Fig. 2e). By contrast, when a mutation changed the amino acid composition or disrupted the reading frame, the model exhibited a much stronger rejection signal. The median ΔPPL associated with missense mutations (0.291) was approximately four-fold higher than that of synonymous mutations (0.076), indicating that the model can distinguish simple nucleotide substitutions from coding changes that are more likely to affect the protein product. For more disruptive mutation types, the increase in perplexity was even more pronounced. Nonsense mutations introducing premature stop codons resulted in a substantial median ΔPPL of 1.732. Stop-loss mutations disrupting the original termination codon also showed comparable increases (median ΔPPL = 1.558), while single-base deletions causing frameshifts produced the strongest overall effect by a wide margin (median ΔPPL = 5.411) (Fig. 2e,f). This overall trend is highly consistent with the degree to which coding-region integrity is compromised.

These results show that, even without any task-specific fine-tuning on variant data, GenNA displays discriminative responses to different classes of mutations solely as a result of large-scale pretraining, consistent with findings in other genomic foundation models^6^. The ΔPPL used here reflects changes in model likelihood induced by perturbations that deviate from the learned distribution of natural sequences, and is therefore treated as a quantitative proxy for mutation-induced deviations in sequence likelihood rather than a direct experimental measurement of phenotypic effects. Nevertheless, the emergence of this stratified pattern in the absence of population-level variation data from the same species suggests that cross-species pretraining can endow the model with sensitivity to coding constraints and local sequence abnormalities.

### Capturing evolutionary constraints and phylogenetic representations

To investigate whether GenNA has learned deeper statistical features associated with species context, we systematically analyzed its intermediate-layer representations and its conditional sensitivity to perturbations in species labels. Although the pretraining corpus explicitly provides only shallow species labels, such as genus and species epithets, rather than a complete hierarchical taxonomy, the model may still implicitly acquire a degree of species-similarity structure by modeling differences among homologous sequences across species^30^.

In the absence of any explicit taxonomic supervision, we extracted hidden states from an intermediate Transformer layer (layer 18) and visualized sequence representations from different species using t-SNE^31^. The resulting embeddings were not randomly dispersed, but instead showed partially separable clustering patterns according to broad biological groups, such as fungi, plants, invertebrates, and vertebrates. In addition, the model formed relatively well-defined subspaces for the two molecular modalities, DNA and RNA (Fig. 2g), indicating that GenNA distinguishes not only between molecular modalities but also differences associated with species context.

We next quantified the model’s sensitivity to species specificity by measuring the ΔPPL induced by incorrect species prompts. We hypothesized that, if the model captures species-associated sequence features, then an incorrect species label should not behave as uninformative noise; rather, it should conflict with the underlying sequence patterns and thereby produce a detectable perturbation in the probability assigned to the entire conditioned input. Using the RNA sequence of the ppap2d gene from the great blue-spotted mudskipper (*Boleophthalmus pectinirostris*) as an example, we found that replacing the original species prompt with labels from highly divergent species, such as human or Arabidopsis thaliana, led to a clear increase in average PPL (Fig. 2h).

To assess the generality of this phenomenon, we further selected 10 representative species and calculated the average ΔPPL produced by all pairwise species-label substitutions, thereby constructing a ΔPPL matrix. This matrix displayed a relatively clear diagonal-consistent structure: in general, the greater the difference between two species, the larger the ΔPPL induced by an incorrect label (Fig. 2i). Hierarchical clustering based on this ΔPPL matrix produced a topology that showed broad agreement with reference phylogenetic relationships across several major branches, including a reasonable separation of fishes, birds, and mammals within vertebrates (Fig. 2j). However, some deviations remained for more distantly related taxa; for example, in the learned representation space, the model tended to place Saccharomyces cerevisiae closer to Chlamydomonas reinhardtii. Overall, these results suggest that what GenNA learns is better characterized as a phylogeny-related structure of statistical similarity rather than as a strict reconstruction of evolutionary relationships^32^.

### Broad and meaningful sequence generation

One of the central goals of GenNA is to enable conditional nucleotide generation guided by natural-language annotations. To evaluate whether the generated outputs are not only formally plausible but also consistent with the underlying biological context in both semantic and statistical terms, we systematically analyzed model behavior across multiple generation settings (Fig. 3a).

**Fig. 3:**
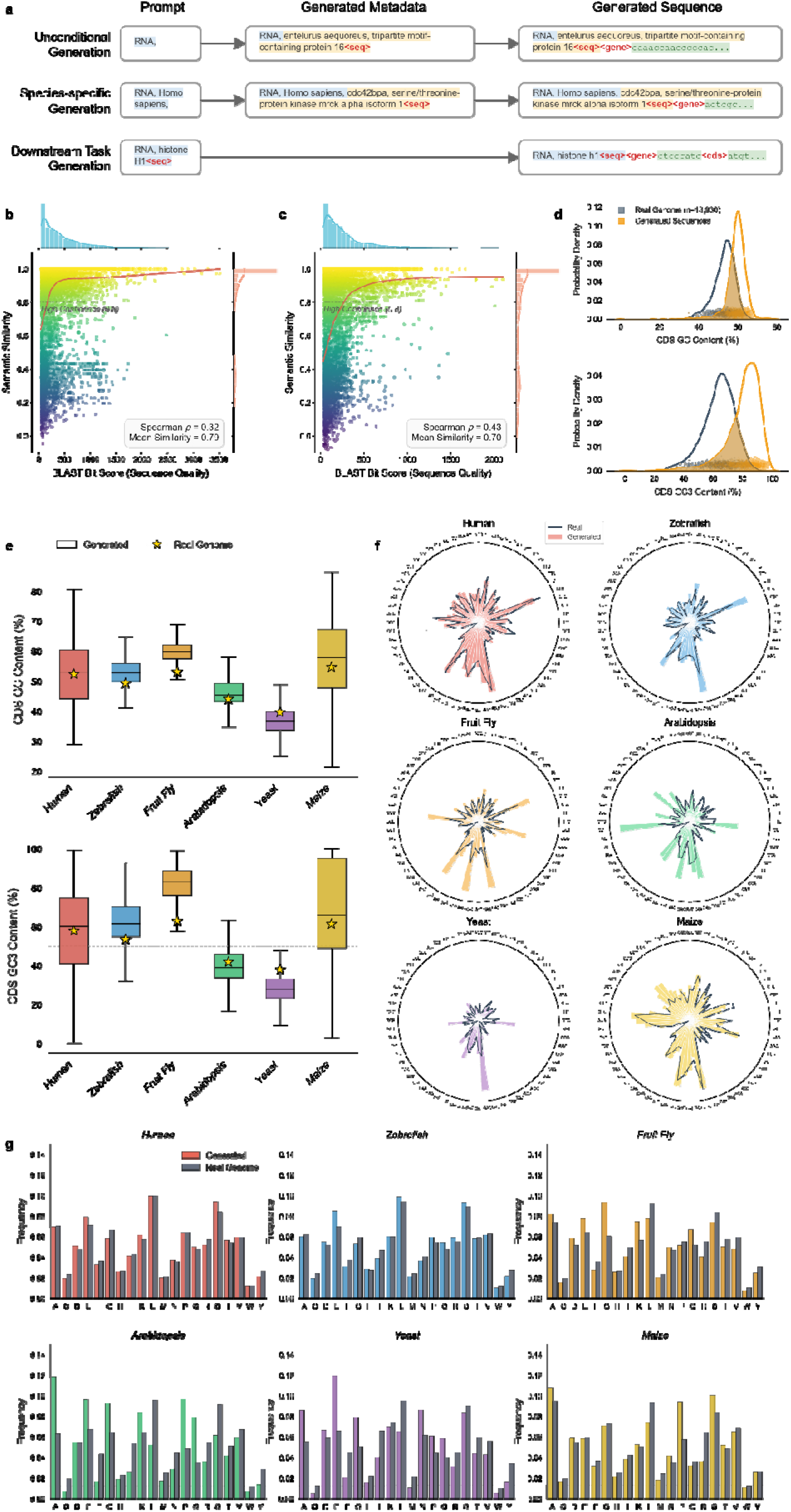
Large-scale unconditional and species-specific sequence generation. **a**, Schematic of the three different generation paradigms: unconditional self-guided generation, species-specific generation, and downstream task-directed generation. **b, c**, Cross-modal semantic consistency of generated sequences from unconditional and species-specific generation tasks. Scatter plots show cosine similarity between generated text metadata and BLASTP-derived functional annotations, with horizontal dashed lines indicating the high-confidence threshold (0.8). Marginal plots show data density. **d**, Probability density distributions of CDS GC content (top) and CDS GC3 content (bottom) for sequences generated by GenNA versus real gene sequences of fruit fly *(Drosophila melanogaster)*. **e**, Comparison of CDS GC (top) and CDS GC3 (bottom) content across six representative eukaryotic model organisms. **f**, Radar charts comparing synonymous codon usage bias between real reference sequences (dashed black lines) and generated sequences (solid colored lines). **g**, Global amino acid frequencies of real and generated sequences across 6 model organisms.

We first evaluated unconditional generation under highly minimal conditions. In this setting, all explicit constraints were removed, and the model was provided only with a modality tag (Genomic DNA or RNA). It was then required to autoregressively generate the complete metadata fields, including species, gene name, and functional annotation, followed by the corresponding nucleotide sequence. For generated mRNA sequences, we extracted the CDS regions and translated them into protein sequences, which were then aligned against existing databases using BLASTP^33^. We subsequently used Sentence-Transformers^34^ to evaluate the semantic consistency between the model-generated functional annotations and the annotations associated with the top database hits. Sequence-logo analysis of these unconditionally generated mRNAs further recovered a Kozak-like translation-initiation context, which is a meaningful local sequence grammar (Supplementary Fig. 5). The results showed that the generated annotations remained well aligned with the corresponding generated sequences at the semantic level (Fig. 3b).

We next evaluated species-specific generation to investigate whether the model could leverage species-specific properties to guide sequence generation. Without imposing any explicit constraint on gene function, we prompted the model to generate nucleotide sequences from six representative model organisms: human *(Homo sapiens)*, zebrafish *(Danio rerio)*, fruit fly *(Drosophila melanogaster), Arabidopsis thaliana*, yeast *(Saccharomyces cerevisiae)*, and maize *(Zea mays)*. In addition to preserving semantic self-consistency (Fig. 3c), GenNA successfully reproduced species-specific features in the generated sequences (Fig. 3d-g). For example, the GC-content distributions of the generated sequences were broadly consistent with those of the corresponding reference genomes, and the model captured systematic differences across species ranging from relatively low-GC to high-GC backgrounds. Similar trends were also observed for GC3, codon-usage bias, and amino acid frequency.

In addition, we observed that the generated distributions were more concentrated than the corresponding natural reference distributions, exhibiting a degree of distribution sharpening. In fruit fly and maize, for example, the generated sequences tended to cluster more strongly around the most representative GC ranges of each species, whereas coverage of the long-tail regions in the natural distributions was relatively limited (Fig. 3e). This pattern suggests that, under the current sampling configuration, GenNA preferentially generates typical samples that conform to the dominant modes of the training distribution while still underrepresenting low-frequency tail variation^35,36^. Overall, these results indicate that GenNA can generate sequences that are not only consistent with its self-generated functional annotations but also preserve species-specific statistical features.

### Controllable generation of nucleotide sequences

To further assess GenNA’s capacity for controllable generation, we performed natural-language-guided targeted generation experiments on non-coding RNAs with structural constraints (tRNAs and rRNAs), as well as on protein-coding genes whose translated products exhibit family-specific physicochemical properties (histones). The model was provided the prompt specifying the generation target and was required to generate the corresponding nucleotide sequence (Fig. 3a).

For tRNAs, we generated a total of 30,500 candidate sequences from prompts covering 61 distinct anticodons and evaluated them systematically using tRNAscan-SE^37^. The results showed that most generated sequences achieved relatively high Cove scores, indicating broad consistency with the secondary-structure constraints of natural tRNAs (Fig. 4a). Further analysis revealed that, when generating tRNAs for specific amino acids, the model spontaneously adopted multiple anticodon-recognition strategies consistent with natural systems, including wobble pairing^38^ (Fig. 4b). In addition, the model reproduced the relatively longer sequence characteristics of tRNA-Leu and tRNA-Ser, which possess a long variable arm (Type II tRNA)^39^ (Fig. 4c). For rRNAs, we generated four classes of sequences, 5S, 5.8S, 18S, and 28S, and aligned them to Rfam covariance models^40^ using cmsearch from the Infernal suite^41^. The resulting bit-score distributions were overall close to those of real rRNA reference sets (Fig. 4d), suggesting that the long-chain rRNAs generated by the model are probabilistically compatible with natural family-level structural models.

**Fig. 4:**
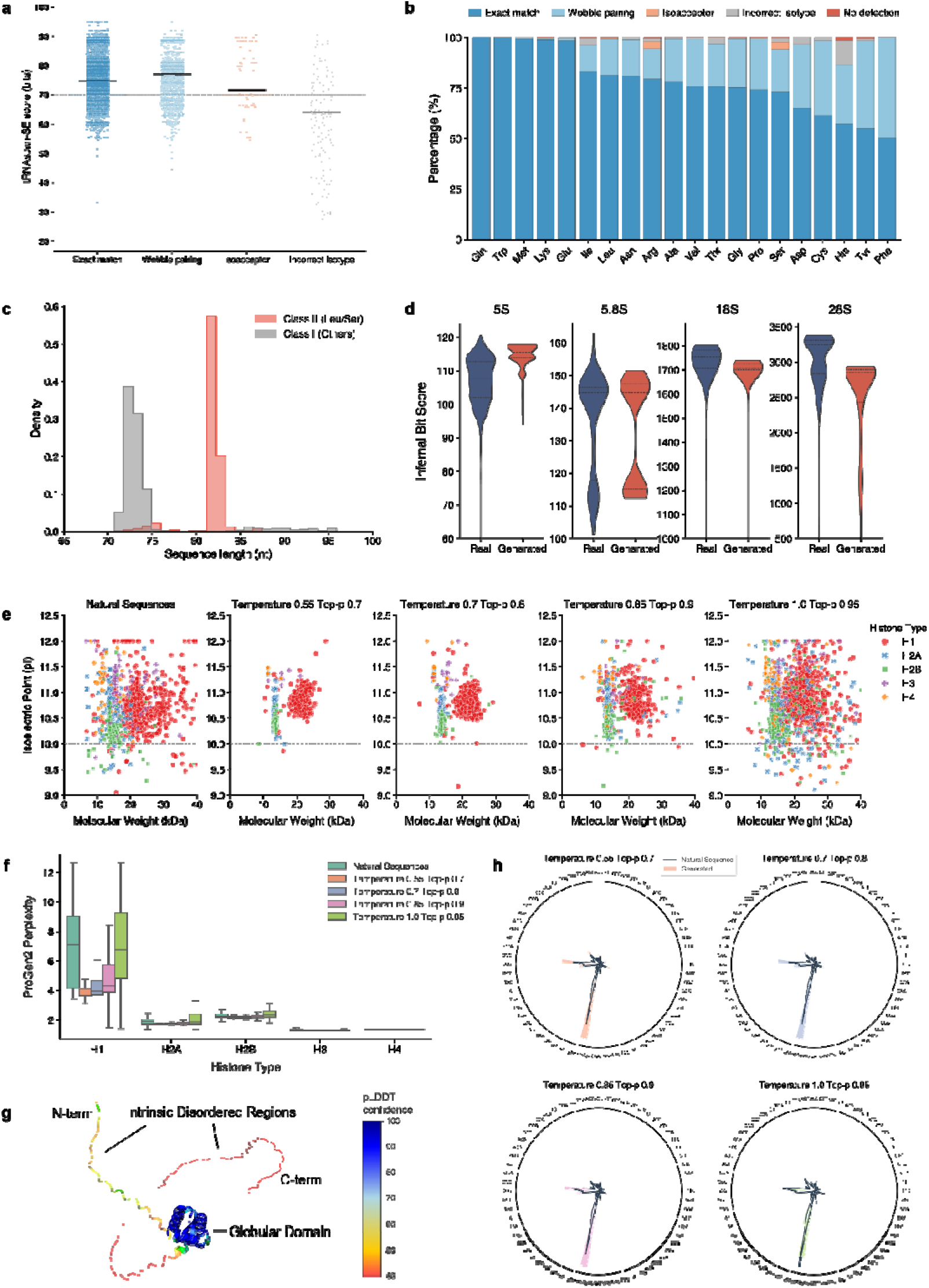
Controllable generation of structured non-coding RNAs and protein-coding sequences. **a**, Distribution of tRNAscan-SE Cove scores for generated tRNAs, categorized by anticodon pairing behavior. Solid horizontal lines represent medians. **b**, Proportion of anticodon pairing strategies spontaneously adopted by the model when generating tRNAs for specific amino acids. **c**, Sequence length distribution for generated Class II (Leu/Ser) and Class I (others) tRNAs. **d**, Infernal bit scores for natural and generated ribosomal RNAs (5S, 5.8S, 18S, 28S). **e**, Physicochemical landscape (isoelectric point and molecular weight) of natural core histones and GenNA-generated histones across incrementally increasing sampling temperatures and top-p thresholds. **f**, ProGen2 perplexity scores for natural and generated sequences across five histone types. **g**, Schematic of the 3D spatial domains of a histone, highlighting intrinsic disordered regions (IDRs). The structure is colored by AlphaFold2 pLDDT confidence score (blue indicates high confidence; warm colors indicate low confidence). **h**, Radar charts showing amino acid composition preferences of natural and generated histones modulated by different sampling parameters.

For protein-coding genes, we further generated sequences corresponding to five major histone classes, H1, H2A, H2B, H3, and H4, and analyzed the distributions of their translated products in molecular-weight (MW) and isoelectric-point (pI) space. The results showed that GenNA was able to recapitulate the principal distribution patterns of different histone families in physicochemical-property space (Fig. 4e). For example, the generated H1 sequences generally retained the trends of higher molecular weight and stronger basicity^42^, whereas generated H3 and H4 samples remained more tightly clustered. Together with the ProGen2 perplexity evaluation^43^ (Fig. 4f), these results indicate that the generated coding sequences preserve biophysical profiles similar to those of natural histone families.

Sampling parameters had a substantial influence on the generated distributions. Using H1 as an example, under lower-temperature and more conservative sampling settings, the generated sequences exhibited a more concentrated, spike-like distribution in features such as length and GC content. As the temperature increased and the sampling threshold was relaxed, the coverage of generated samples in physicochemical-property space expanded markedly, and the resulting continuous distributions became closer to those of the natural reference set (Fig. 4e,h and Supplementary Fig. 6). Overall, these results suggest that GenNA captures multilevel biological features ranging from RNA secondary structure to protein physicochemical properties, while also exhibiting a degree of controllability at inference time through autoregressive sampling parameters. By adjusting prompts and sampling settings, the model can switch between more conservative targeted generation and more open-ended sequence exploration.

## Discussion

In this study, we introduced GenNA, a generative large language model for nucleotide sequences guided by natural-language annotations. Unlike prior approaches that are primarily designed to model a single sequence modality, GenNA employs a dedicated cross-modal BPE vocabulary and joint pretraining on large-scale eukaryotic genomic and transcriptomic corpora to establish associations between functional semantics and sequence syntax within a unified autoregressive framework. Our results show that this training strategy not only enables the model to capture the intrinsic patterns of nucleotide sequences themselves, but also allows it to leverage natural-language functional annotations and species context for consistency evaluation and conditional generation over entire conditioned inputs.

GenNA exhibits several notable emergent properties at multiple levels. First, in in silico mutational scanning, the model shows a stratified perplexity response across synonymous, missense, nonsense, and frameshift mutations, with more disruptive variants generally associated with larger ΔPPL. This pattern suggests that the model has learned statistical features related to genetic-code degeneracy and coding-sequence integrity. Second, even without explicit supervision from a taxonomic tree, both the intermediate-layer representations of GenNA and the ΔPPL matrix derived from species-label swapping reveal a degree of species-associated structure, indicating that cross-species pretraining may enable the model to capture phylogeny-related latent similarities^30^. Furthermore, in the tRNA generation task, the model spontaneously reproduces wobble-pairing behavior consistent with natural systems, suggesting that cross-modal training may influence not only local sequence continuation but also the internalization of higher-order biological constraints.

Taken together, these findings support a broader conclusion: for nucleotide foundation models, introducing natural-language annotations as explicit conditioning inputs not only improves the model’s ability to exploit sequence-function pairings, but also provides a more natural interface for generating candidate sequences from high-level biological intentions. At the same time, it is important to emphasize that many of the evaluations in this study are still based primarily on computational proxy metrics, including PPL over entire conditioned inputs, perplexity from external models, covariance-model matching scores, and distributions of physicochemical properties. Accordingly, these results are better interpreted as evidence that the model has learned probabilistic constraints and structural compatibilities consistent with natural biological sequences, rather than as direct confirmation of true biological function, experimental phenotype, or in vivo feasibility.

Despite the encouraging performance of GenNA across multiple tasks, several limitations remain. First, due to computational constraints, we have not yet systematically explored how scaling laws affect the upper bound of model performance^44^. Second, in sequence-generation tasks, we observed that the generated distributions are often more concentrated than those of real natural genomes; that is, they exhibit distribution sharpening. This behavior may be related to truncation of low-probability tail regions by top-p sampling during autoregressive decoding^35^, and it suggests that more refined sampling-calibration strategies ought to be explored in future work to better recover the diversity present in natural sequence distributions.

In addition, there remains room for improvement in both tokenization strategy and context length. Although the 6,000-token BPE vocabulary used here provides a reasonable trade-off between sequence compression and the capture of biological motifs, it may still lose part of the local information required in settings that demand single-nucleotide resolution, such as highly fine-grained mutational analyses of cis-regulatory elements. Future work could explore hierarchical tokenization, mixed-granularity modeling, or longer-context architectures to improve the joint modeling of ultralong-range regulatory relationships and fine-scale local variation. Another important limitation is that all generated results in this study currently lack systematic wet-lab validation. Consequently, whether in terms of the structural compatibility of generated tRNAs/rRNAs or the plausibility of histone-coding sequences, further experimental verification will still be required.

Looking ahead, we plan to further extend the applicability of GenNA to more complex downstream tasks through parameter-efficient fine-tuning (PEFT), such as LoRA^45^. Within such a framework, GenNA could potentially serve both as a conditional sequence generator and as a zero-shot or few-shot discriminator, supporting tasks such as candidate-sequence design in synthetic biology and variant prioritization. As training data continue to expand, particularly through the incorporation of more functional assays, population-variation data, and multi-omics annotations, cross-modal nucleotide foundation models such as GenNA may provide an increasingly tight link between biological sequence understanding and sequence design.

## Methods

### Pretraining corpus construction

To construct the multimodal nucleotide pretraining corpus, we collected all available GenBank-formatted^26^ genomic DNA and RNA annotation files for eukaryotic species from the NCBI RefSeq database^25^. Because intergenic and other non-gene regions often contain abundant repetitive and low-complexity sequence, and a genomic language-modeling study reported better downstream performance when training was concentrated on annotated gene regions rather than mixed whole-genome sequence^17^, we primarily restricted the corpus to gene-associated regions with explicit annotations.

We used Biopython^46^ to parse GenBank records automatically. For each valid record, we extracted the molecule type (genomic DNA or RNA), source species, gene name, and natural-language functional descriptions from the product and note fields. For hierarchical structures such as mRNA, CDS, ncRNA, and regulatory elements, we organized the annotations using custom XML-style tags so as to preserve the original structural boundaries and semantic cues as much as possible, and wrote these structured annotations together with the corresponding underlying nucleotide sequences into unified training samples.

During construction of genomic DNA samples, to introduce limited local neighborhood context while preserving the core annotated region, we randomly extended the target annotated segment by 0–100 bp upstream and downstream; extensions were truncated when they reached the physical boundary of the corresponding contig. After cleaning and reorganization, the final corpus comprised approximately 24.7 million samples and approximately 416 billion characters in total. To improve context utilization and GPU memory efficiency during Transformer pretraining, we further applied a sequence-packing strategy during preprocessing, concatenating multiple samples into long input sequences of nearly 20,000 characters^47^.

### Cross-modal BPE tokenizer

Because the pretraining corpus simultaneously contained continuous nucleotide sequences, natural-language functional annotations, and XML-style structural tags, the model required a tokenization strategy capable of handling multiple symbol types in a unified manner. Traditional nucleotide k-mer tokenization is suitable for pure sequence modeling but cannot readily encode natural-language phrases or hierarchical tags^5,6^. By contrast, character-level encoding is more general but substantially increases input token length, thereby reducing the computational efficiency of long-context training^48^.

On the basis of these considerations, we used byte-pair encoding (BPE)^23^ to construct a unified cross-modal tokenizer. We randomly sampled 10,000 records from the pretraining corpus as the tokenizer training set and trained a dedicated vocabulary of 6,000 tokens. This vocabulary size provided a balance between local nucleotide-pattern resolution and overall compression efficiency: it preserved sensitivity to short nucleotide fragments and local variation, while compressing high-frequency natural-language functional phrases, common XML tag fragments, and recurrent sequence motifs into more stable discrete units.

During unsupervised BPE merging, high-frequency co-occurrence patterns from different modalities were mapped into a shared token space. This design enabled the model to process sequence syntax, textual semantics, and structural boundary information jointly within a common input representation, thereby providing a unified foundation for subsequent causal language modeling.

### Model architecture and configuration

GenNA used a decoder-only Transformer architecture. Its core network configuration was based on the Qwen3-4B design paradigm^22^, but the model was reinitialized and pretrained from scratch for the present cross-modal nucleotide modeling task. Because we used a custom cross-modal BPE vocabulary rather than the original Qwen3 vocabulary, the model parameters were not transferred from existing checkpoints; instead, the model underwent full pretraining under the new vocabulary system and multimodal nucleotide corpus.

The model comprised 36 Transformer decoder blocks with a hidden size of 2,560. The attention module used grouped query attention (GQA)^49^, with 32 query heads and 8 key/value heads. Rotary positional embeddings (RoPE)^50^ were used for positional encoding, the feed-forward network employed the SwiGLU activation function^51^, and layer normalization used RMSNorm^52^ in a pre-norm configuration.

Under this configuration and the 6,000-token vocabulary, the model contained approximately 3.6 billion parameters in total. This architecture was designed to maintain strong long-range modeling capability together with reasonable training and inference efficiency, while accommodating inputs that combined nucleotide sequences, natural-language prompts, and structured tags.

### Pretraining protocol

Pretraining was carried out on a computing cluster of five NVIDIA A800 GPUs (80 GB memory each). The training objective was causal language modeling (CLM), i.e., prediction of the next token conditioned on the preceding context^20^. For an input token sequence *x* = (*x*_1_, *x*_2_, …, *x*_*T*_) of length *T*, the objective can be written as:

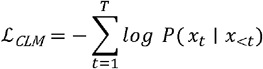

where *P*(*x*_*t*_ | *x*_<*t*_) denotes the conditional probability assigned by the model to the *t*-th token given the preceding tokens.

The optimizer was AdamW^53^, with *β*_1_ = 0.9, *β*_2_ = 0.95, and *λ*_*weight decay*_ = 0.1. The learning-rate schedule combined linear warmup with cosine annealing^54^. During the first 5,000 update steps, the learning rate increased linearly from 0 to 2 × 10^−4^; it then decayed gradually according to a cosine schedule to 10% of the peak value over the remainder of training. With gradient accumulation, the effective global batch size was stabilized at 60. Training ran for 400,000 steps in total and required approximately 3 months of computation.

To reduce the model’s tendency to rely mechanically on specific explicit prompt items, such as Latin species names or fixed gene symbols, and to increase its dependence on more general contextual statistical patterns, we introduced a dynamic information-masking strategy the during the later stage of training. During construction of training inputs, the species identifier or gene identifier in the prompt was randomly removed with probability 0.5.

### Evaluation of embeddings and inference metrics

Within the evaluation framework of this study, perplexity (PPL) was used as the core metric for quantifying the model’s overall uncertainty with respect to an input sample. For an input token sequence *X* = (*x*_1_, *x*_2_, …, *x*_*T*_) formed by the prompt and the sequence under evaluation, the average perplexity was defined as:

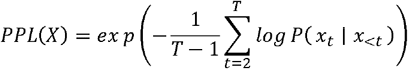

This definition is equivalent to the exponential transform of the average negative log-likelihood over the entire input sequence:

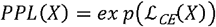

where

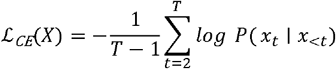

is the mean cross-entropy loss over the full input token sequence.

### In silico mutation scanning

To evaluate the model’s sensitivity to local sequence perturbations systematically, we established an in-silico mutation pipeline based on protein-coding genes from the validation set. We randomly selected 100 protein-coding genes with lengths between 200 and 3,000 bp from an independent validation set as the wild-type sequence collection (*S*_*wt*_). For each gene, we exhaustively introduced single-nucleotide substitutions and single-base deletions at each position in the untranslated regions (5′ UTR and 3′ UTR) and coding sequence (CDS).

According to the standard genetic code for nuclear protein-coding genes, simulated mutations in the CDS were further classified as synonymous, missense, nonsense, and stop-loss mutations^55^. Single-base deletions occurring within the CDS that altered the reading-frame phase were classified separately as frameshift mutations. The perturbation caused by each mutation was quantified by the perplexity difference between the mutant sequence (*S*_*mut*_) and the wild-type sequence (*S*_wt_):

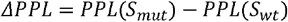

A positive ΔPPL indicated that the model regarded the mutant sequence as further removed from the natural distribution it had learned than the corresponding wild-type sequence. We therefore used ΔPPL as a proxy for the degree of abnormality and to compare the relative magnitude of perturbations induced by different mutation classes^28^.

### Sequence-function semantic consistency evaluation

To evaluate the model’s cross-modal alignment between natural-language functional descriptions and nucleotide sequences, we constructed a validation set containing *N* real gene samples, where each sample consisted of a nucleotide sequence *S*_*i*_ and its corresponding native functional annotation *A*_*i*_. In this experiment, the underlying nucleotide sequence was held fixed, while the functional-annotation component of the prompt was systematically replaced, thereby generating inputs with either matched or mismatched sequence-function pairings.

Specifically, for each sequence *S*_*i*_ in the collection, we combined it with every annotation *A*_*j*_ to form input prompts and computed the corresponding conditional perplexity. The resulting evaluations were divided into two groups: the matched group (*i* = *j*) in which each sequence was paired with its original annotation, and the mismatched group (*i* ≠ *j*), in which the annotation was replaced. These comparisons were used to assess the direct influence of semantic constraints on the probability assigned to nucleotide sequences.

### Model embedding and phylogenetic representation analysis

To determine whether GenNA internalizes species-specific signatures, we extracted continuous hidden representations from intermediate layers of the pretrained model. Specifically, we used standardized prompts of the form `“Genomic DNA, [Species Name]<seq>“` or `“RNA, [Species Name]<seq>“` and captured the hidden states from layer 18 of the 36-layer Transformer decoder. Mean pooling was applied across the sequence dimension, excluding padding tokens, to obtain a fixed-dimensional global embedding for each sequence. These embeddings were visualized using t-distributed stochastic neighbor embedding (t-SNE)^31^, with perplexity = 30 and maximum iterations = 1,000.

We further assessed the model’s conditional sensitivity to changes in species labels using PPL perturbation. Ten representative species spanning the major eukaryotic lineages were selected. For any species pair (*i, j*), the species label in the prompt was replaced with that of species *j*, while the rest of the input was kept unchanged, and the target sequence from species *i* was then evaluated. The corresponding average ΔPPL was defined as:

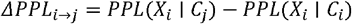

where *X*_*i*_ denotes the target sequence from species *i*, and *C*_*i*_ and *C*_*j*_ denote the conditioning prompts constructed with the correct and substituted species labels, respectively. The resulting 10 × 10 ΔPPL matrix was subjected to hierarchical clustering using Ward’s minimum-variance method^56^, and the resulting topology was compared with a reference phylogenetic tree^57^.

### Sequence generation and validation

#### Autoregressive sequence generation framework and decoding strategy

To enable conditional de novo generation, GenNA used a unified prompt template as the input prefix:

~~~
[Molecule Type], [Species Name], [Gene Symbol], [Functional Annotation]<seq>
~~~

where the species name and gene symbol could be optionally provided depending on the task. Given a prefix C, the model autoregressively generated a target token sequence *Y* = (*y*_1_, *y*_2_, …, *y*_*T*_), with joint conditional probability:

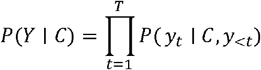

At each step, the model first produced a logits vector *z*_*t*_, which was converted to a probability distribution by softmax. To control the smoothness of the sampling distribution, temperature scaling was applied during inference:

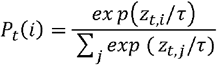

where *τ* is the temperature parameter. When *τ* < 1, the distribution becomes sharper and sampling is biased toward higher-probability tokens; when *τ* > 1, the distribution becomes flatter and the generated outputs are typically more diverse.

After temperature scaling, nucleus sampling (top-p sampling) was applied^35^. Specifically, token probabilities at the current step were sorted in descending order, and the smallest candidate set *V*_*p*_ satisfying a cumulative probability of at least was selected:

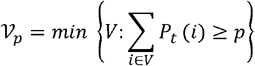

Sampling was then performed only within this candidate set. To suppress repetitive fragments, a repetition penalty was applied to previously generated tokens, and repeated n-grams of length were disallowed. Unless otherwise specified, the following decoding parameters were used throughout: temperature = 0.7, top-p = 0.8, repetition penalty = 1.3^58^, and no-repeat n-gram size = 5. Generation ended when the model emitted the terminator `<eos>` or reached the predefined maximum token length.

#### Unconditional self-generation and semantic fidelity analysis

In the unconditional self-guided generation task, the model received only the modality label (Genomic DNA or RNA) as the initial input and autoregressively generated the complete metadata fields, including species, gene name, functional annotation, and the subsequent nucleotide sequence. We generated a total of 10,000 samples.

To approximate the consistency between generated sequences and their self-generated annotations at the protein level, we extracted the CDS region enclosed by the `<cds></cds>` tags from each generated mRNA sequence and translated it into an amino acid sequence. These translated sequences were then aligned against the UniProt database^59^ using BLASTP^33^, and the functional annotations associated with the top five hits ranked by bit score were retained. We further used a pretrained semantic model, Sentence-Transformers (all-MiniLM-L6-v2)^60^, to compute the cosine similarity between the model-generated annotation and the BLASTP-hit annotations as an approximate measure of semantic consistency^34^.

#### Species-specific genome feature evaluation

For the species-specific generation task, we selected six model eukaryotes as conditioning species: *Homo sapiens, Danio rerio, Drosophila melanogaster, Arabidopsis thaliana, Zea mays*, and *Saccharomyces cerevisiae*. Without explicitly constraining gene function, we generated 2,000 RNA sequences for each species.

In addition to using the same annotation-consistency analysis as in the unconditional generation experiment, we extracted the CDS regions from the generated mRNA sequences and compared them with species-specific reference CDS statistics from the Codon Statistics Database^61^. The evaluation metrics included the global GC-content distribution, GC content at the third codon position (GC3), codon-usage bias, and amino acid usage frequency. These statistics were used to assess whether the model could reproduce species-associated compositional features during conditional generation.

#### tRNA generation

We conducted targeted generation experiments for transfer RNAs (tRNAs). For 61 tRNA prompt categories with different anti-codon, we constructed inputs using the format `“Genomic DNA, [tRNA Name]<seq>“`, for example, `“Genomic DNA, trnaA-AGC, trna-Ala<seq>“`. Under each prompt condition, 500 samples were generated, yielding a total of 30,500 candidate sequences.

The generated sequences were evaluated using tRNAscan-SE v2.0^37^ with the default eukaryotic setting (-E). The reported Cove score was recorded as an indicator of secondary-structure plausibility. We additionally parsed the anticodon and isotype annotations returned by the tool to assess category-level consistency between the generated sequences and the target tRNA classes.

#### rRNA generation

We evaluated the model’s ability to generate different classes of ribosomal RNAs (rRNAs), including 5S, 5.8S, 18S, and 28S rRNAs. Inputs were constructed using the template `“RNA, [rRNA Name]<seq>“`, for example, `“RNA, 18S ribosomal RNA<seq>“`. For each rRNA class, 1,000 candidate sequences were generated. In parallel, we extracted real rRNA sequences of the corresponding classes from RefSeq eukaryotic GenBank annotation files to construct natural reference sets for comparison.

To evaluate the consistency of the generated sequences with natural rRNA family structural models, we aligned the generated sequences against the corresponding covariance models in the Rfam database^40^ using cmsearch from the Infernal package^41^. The returned alignment scores were used as approximate indicators of structural compatibility. The same procedure was applied to the real rRNA reference sets, allowing comparison between generated and natural sequences in terms of their family-model matching distributions.

#### Histone generation

We used the histone family as a representative task to evaluate the model’s ability to generate coding sequences for specific protein families conditionally. Inputs were constructed using the template “RNA, [histone]<seq>“, where [histone] was specified as H1, H2A, H2B, H3, or H4, corresponding to RNA sequences encoding the respective histone classes. In parallel, we extracted the corresponding real histone-coding sequences, including variants such as H3.1, from RefSeq eukaryotic GenBank annotation files to construct natural reference sets for subsequent comparison.

To examine the effect of sampling hyperparameters on the generated distributions, we used four progressively increasing combinations of temperature and nucleus-sampling threshold: (0.55, 0.7), (0.7, 0.8), (0.85, 0.9), and (1.0, 0.95). In all experiments, the repetition penalty was fixed at 1.3 and the no-repeat n-gram size at 5. For each histone class and each sampling-parameter combination, the model generated 1,000 candidate RNA sequences.

The generated RNA sequences were translated into amino acid sequences to compare with the natural histone sets. Comparison metrics included protein sequence length, molecular weight (MW), theoretical isoelectric point (pI), and amino acid composition, which were computed using the `ProteinAnalysis` class in Biopython’s `Bio.SeqUtils.ProtParam` module^46^. These metrics were used to assess how closely the generated sequences approximated the distributional properties of natural histone proteins.

## Data availability

The nucleotide sequence data used for pretraining were retrieved from the NCBI RefSeq database (https://ftp.ncbi.nlm.nih.gov/genomes/refseq/)^25^. Python scripts used to parse the full-format GenBank files and systematically reconstruct the aligned pretraining corpus are available in our GitHub repository (https://github.com/DrBlackZJU/GenNA).

For downstream validation and sequence analysis, several public databases were utilized: protein sequences for alignment and semantic consistency evaluation were obtained from UniProt (https://www.uniprot.org/)^59^; covariance models for evaluating the structural plausibility of generated rRNAs were sourced from the Rfam database (https://rfam.org/)^40^; reference species-specific CDS statistics, including GC content and codon usage bias, were derived from the Codon Statistics Database (http://codonstatsdb.unr.edu/)^61^. The generated sequences for all evaluated tasks (including unconditional generation, species-specific generation, tRNAs, rRNAs, and histones) are provided in Supplementary Data 1.

## Code availability

The GenNA source code, including scripts for pretraining, generation, and further analysis, is available on GitHub (https://github.com/DrBlackZJU/GenNA). Model weights are publicly available on Hugging Face (https://huggingface.co/DrBlack/GenNA).

## Acknowledgements

This work was supported by the National Natural Science Foundation of China [32270709,32570787,32300532]; the National Key Research and Development Program of China [2023YFE0112300]; the 151 Talent Project, and Science and Technology Innovation Leader of Zhejiang Province [2022R52035]; and Jiangsu Collaborative Innovation Center for Modern Crop Production and Collaborative Innovation Center for Modern Crop Production co-sponsored by province and ministry.

## Author contributions

M.C. and C.F. supervised the study. Y.S. designed the study, collected data, trained the model, and performed further analysis with support from G.C. and J.W. Y.S. wrote the manuscript with input from G.C. and D.C. All authors reviewed and approved the submitted manuscript.

