## Supplementary Information for "GenNA: Conditional generation of nucleotide sequences guided by natural-language annotations"

### Supplementary Table 1: Taxonomic distribution of eukaryotic species in the pretraining corpus

| Species | Count | Percentage |
|---|---:|---:|
| fungi | 665 | 29.9% |
| vertebrate other | 500 | 22.5% |
| invertebrate | 486 | 21.9% |
| vertebrate mammalian | 250 | 11.3% |
| plant | 199 | 9.0% |
| protozoa | 121 | 5.5% |
| Total | 2221 | 100.0% |

Number and percentage of the 2,221 eukaryotic species categorized by major taxonomic groups.

### Supplementary Table 2: Components and serialization rules of the GenNA pretraining corpus

<table>
<tr><th>Component</th><th>Genomic DNA</th><th>RNA</th><th>Output</th></tr>
<tr><td>Molecule type</td><td>Genomic DNA</td><td>RNA</td><td rowspan="4">Header prompt</td></tr>
<tr><td>Species name</td><td colspan="2">Source species</td></tr>
<tr><td>Gene symbol</td><td colspan="2">"gene" qualifier of gene feature</td></tr>
<tr><td>Functional annotation</td><td>"product" of first child feature, or "note" if "product" is unavailable</td><td>"product" of first RNA feature, or "note" if "product" is unavailable</td></tr>
<tr><td>Gene sequence</td><td colspan="2">Genomic or transcript sequence corresponding to the selected gene region</td><td rowspan="2">Nucleotide sequence</td></tr>
<tr><td>Flanking sequence</td><td>Random 0–100 bp upstream and downstream</td><td>Not applied</td></tr>
<tr><td>Gene tag</td><td colspan="2">Gene body wrapped by <gene> ... </gene></td><td rowspan="2">XML-style tags</td></tr>
<tr><td>Internal tags</td><td>Including mRNA, CDS, pseudo, specific ncRNA_class</td><td>Including CDS, tRNA, rRNA, precursor_RNA, sig_peptide, specific ncRNA_class</td></tr>
<tr><td>Whole sequence</td><td colspan="2">[Molecule Type], [Species Name], [Gene Symbol], [Functional Annotation] <seq>...</seq><eos></td><td>Training sample</td></tr>
</table>

**Supplementary Table 3: Cosine similarity of specific k-mer tokens in the embedding layer**

| Rank | Query Token: tggtgaaa | | | Query Token: tgga | | |
|---|---|---|---|---|---|---|
| | Token ID | Token | Cosine Similarity | Token ID | Token | Cosine Similarity |
| 1 | 1250 | tggtgaa | 0.303 | 204 | tggg | 0.326 |
| 2 | 295 | tggtg | 0.250 | 221 | tggc | 0.290 |
| 3 | 3992 | cagtgaaa | 0.249 | 200 | caga | 0.278 |
| 4 | 3308 | tggtaaaa | 0.239 | 197 | tgaa | 0.267 |
| 5 | 644 | tggtga | 0.227 | 192 | caca | 0.220 |
| 6 | 1949 | tggtgac | 0.223 | 5824 | tggagtga | 0.217 |
| 7 | 4823 | tggtgcaa | 0.200 | 202 | cagg | 0.213 |
| 8 | 4919 | tggtggtg | 0.195 | 218 | tgcc | 0.206 |
| 9 | 5463 | tggccaaa | 0.189 | 5349 | tggacaca | 0.203 |
| 10 | 3377 | tggggaaa | 0.184 | 5175 | tggatac | 0.195 |
| 11 | 3391 | tgatgaaa | 0.173 | 5550 | tggacatt | 0.190 |
| 12 | 5862 | tgaaataa | 0.163 | 5705 | tggacagg | 0.185 |
| 13 | 3496 | tggtggg | 0.162 | 5406 | tggagatt | 0.180 |
| 14 | 1978 | tagtgaa | 0.159 | 256 | taga | 0.180 |
| 15 | 3646 | tagtaaaa | 0.158 | 208 | cagc | 0.179 |
| 16 | 2176 | tggagaaa | 0.157 | 5381 | tggagaaaa | 0.178 |
| 17 | 2493 | tggaaaaa | 0.154 | 190 | tgtg | 0.178 |
| 18 | 4424 | tggttcaa | 0.154 | 5528 | tggagagc | 0.176 |
| 19 | 271 | tgaaa | 0.153 | 195 | tgtt | 0.170 |
| 20 | 1853 | tggttaa | 0.150 | 262 | ggga | 0.170 |

Top 20 tokens with the highest cosine similarity to the query sequences 'tggtgaaa' (left) and 'tgga' (right), extracted from the GenNA vocabulary embeddings.

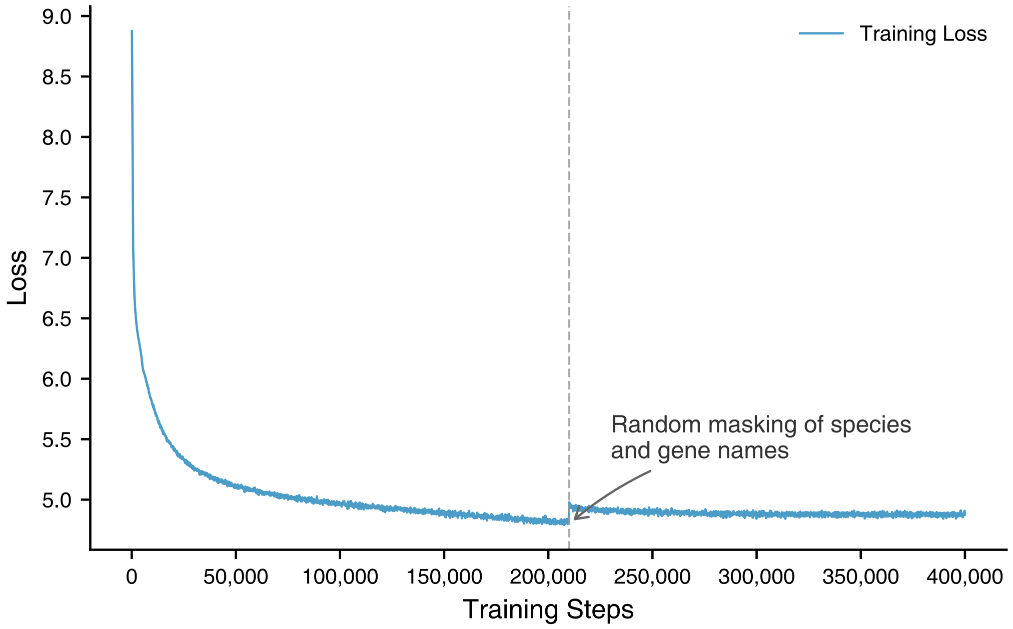

**Supplementary Fig. 1: Pretraining loss curve of the GenNA-small model**

The CLM training loss of the 0.36B parameter version of GenNA over 400,000 training steps. At step 210,000 (indicated by the vertical dashed line), a dynamic information-masking strategy was introduced, under which species and gene identifiers in the prompts were randomly masked with a probability of 0.5.

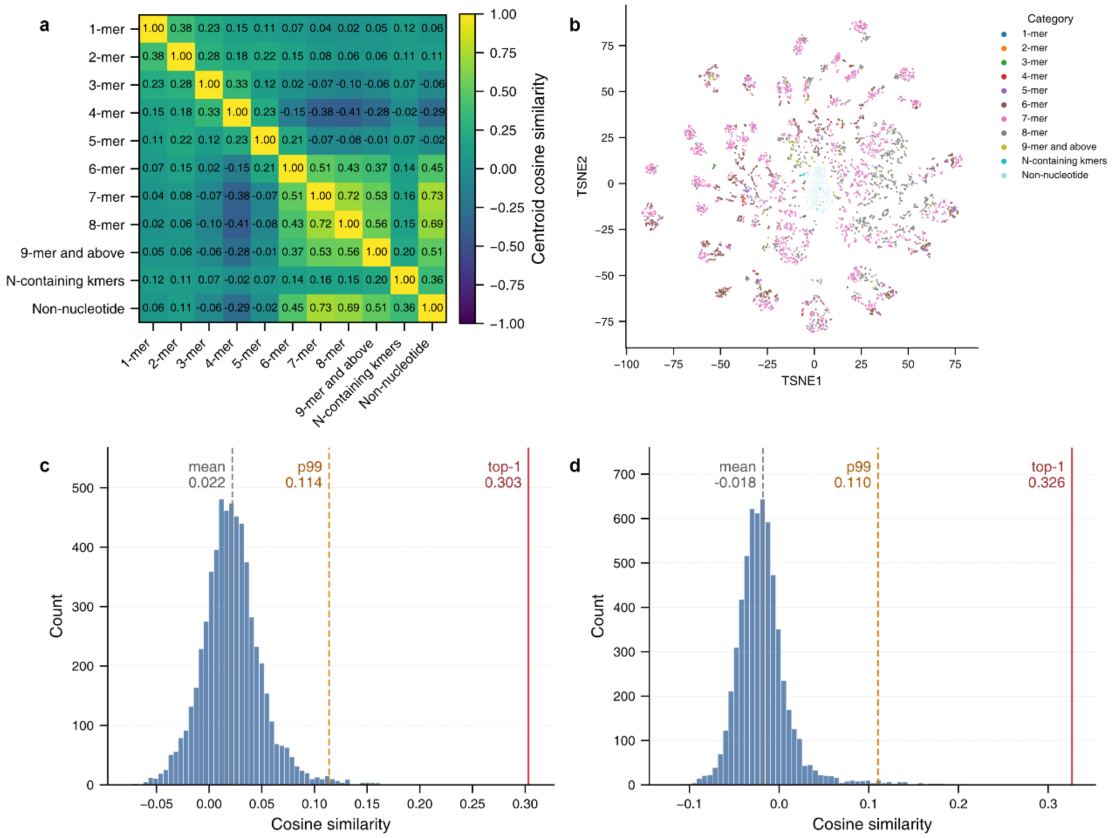

### Supplementary Fig. 2: Visualization of the GenNA vocabulary token embedding space

**a**, Cosine similarity matrix of token category centroids in the embedding layer. Tokens are grouped by nucleotide length (k-mers), presence of ambiguous bases (N-containing), and non-nucleotide status. **b**, Two-dimensional t-SNE[1] projection of the entire 6,000-token vocabulary, colored by category, illustrating the latent clustering patterns based on token composition. **c, d**, Global distribution of cosine similarities across the entire vocabulary relative to the representative query tokens 'tggtgaaa' (**c**) and 'tgga' (**d**).

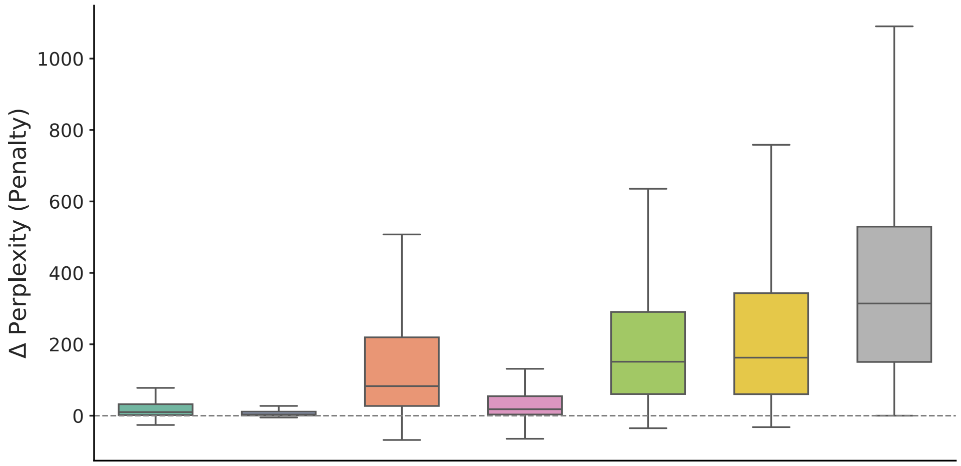

| | | | | | | |
|---|---|---|---|---|---|---|
| Species Name | | ✓ | ✓ | | | ✓ |
| Gene Symbol | ✓ | | ✓ | | ✓ | |
| Functional Annotation | ✓ | ✓ | | ✓ | | |

**Supplementary Fig. 3: Prompt ablation reveals the contributions of different prompt components to GenNA's functional understanding**

ΔPPL distributions after systematic prompt ablation across 5,000 sequences. Prompt components include species name, gene symbol, and functional annotation. Check marks in the table indicate which components are retained in each ablation setting. The prompt ablation results show that functional annotation provides the dominant semantic signal.

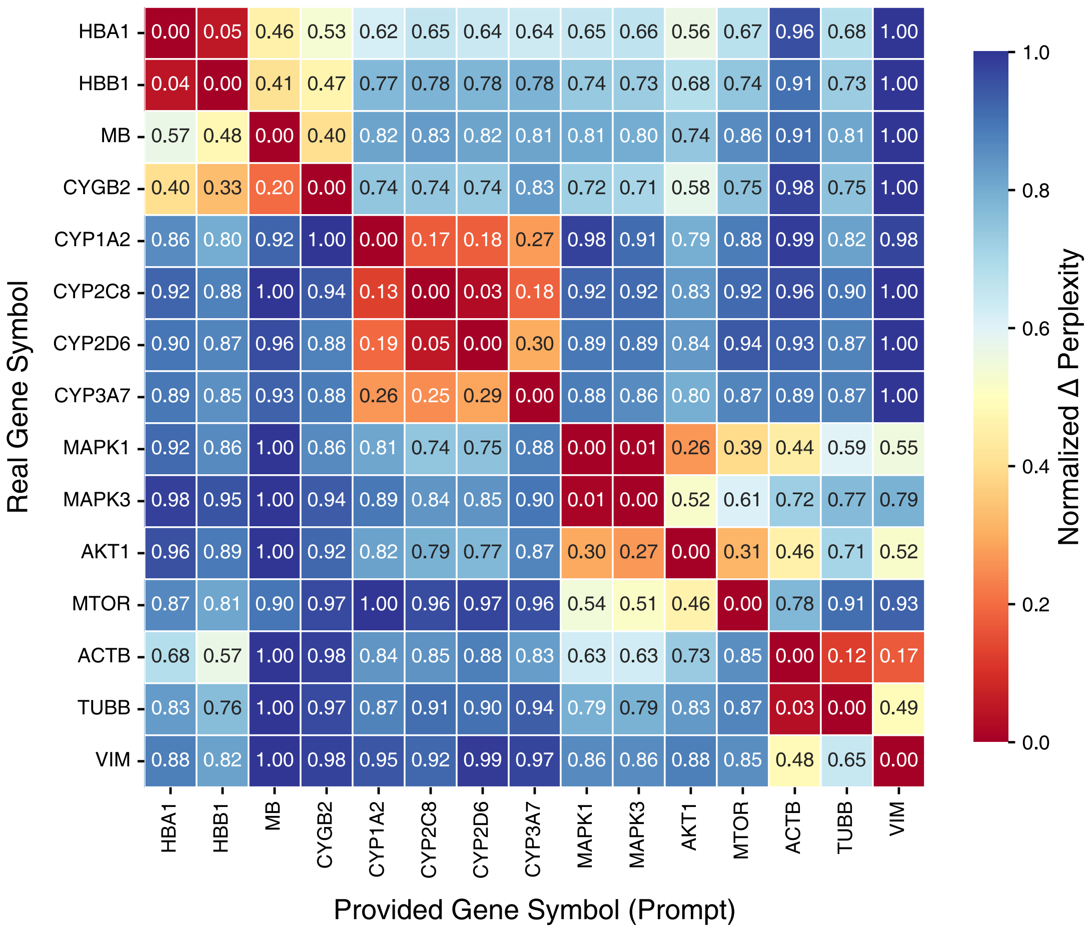

**Supplementary Fig. 4: GenNA distinguishes intra- and inter-family gene similarities**

The normalized ΔPPL heatmap when swapping gene annotations among 15 genes from 4 distinct families. These genes include: globins (HBA1, HBB1, MB, CYGB2), cytochrome P450s (CYP1A2, CYP2C8, CYP2D6, CYP3A7), kinases (MAPK1, MAPK3, AKT1, MTOR), and cytoskeletal components (ACTB, TUBB, VIM).

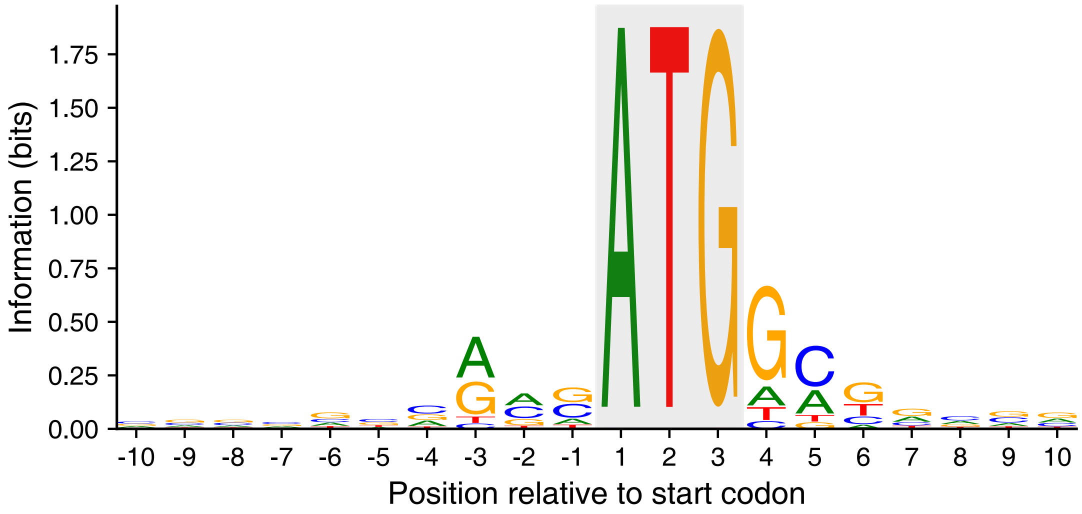

**Supplementary Fig. 5: Sequence logo of the translation initiation context in unconditionally generated mRNA sequences**

The logo[2] displays the information content (bits) of the nucleotide frequency distribution from positions -10 to +10 relative to the start codon (ATG, highlighted in gray) across transcripts produced during the unconditional self-guided generation task. The prominent conservation of a purine (A/G) at the -3 position and a guanine (G) at the +4 position accurately recapitulates the canonical eukaryotic Kozak consensus sequence[3].

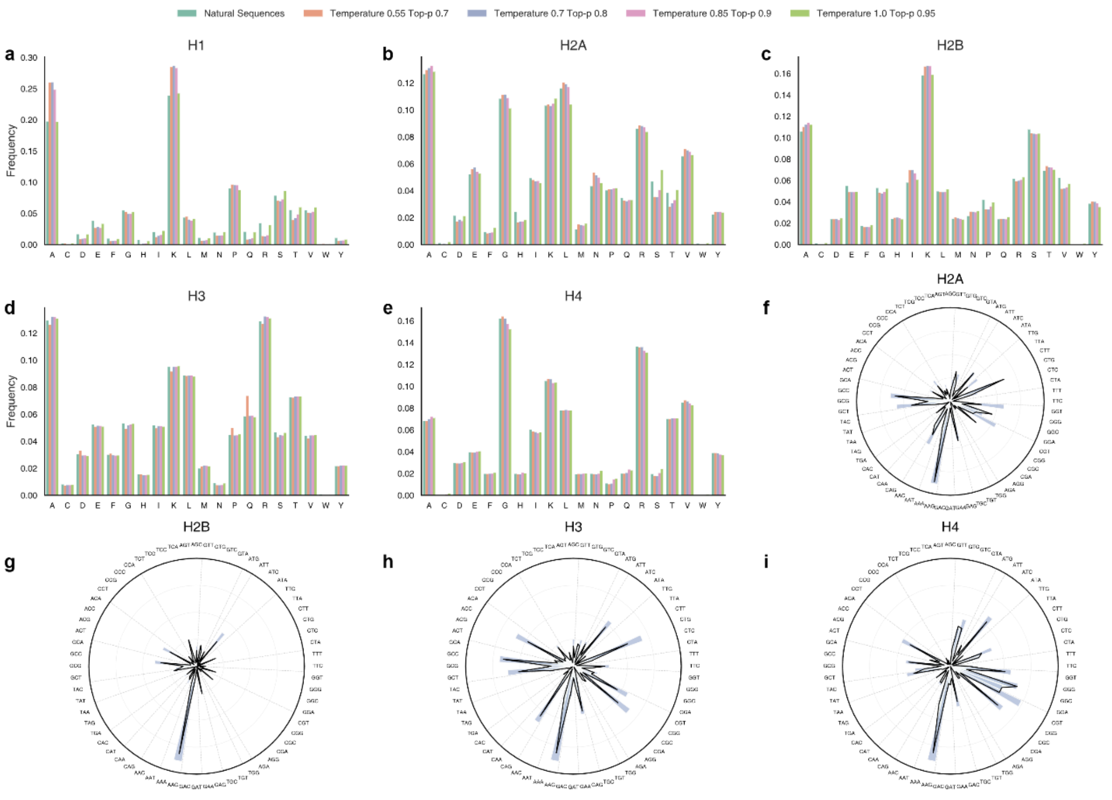

### Supplementary Fig. 6: Family-specific amino acid composition and codon usage preferences of generated histone sequences

**a–e**, The amino acid composition profiles of H1, H2A, H2B, H3, and H4, comparing natural sequences with sequences generated under different sampling hyperparameters. **f–i**, Codon usage preference for H2A, H2B, H3, and H4, comparing natural sequences with sequences generated at temperature 0.7 and top-p 0.8.

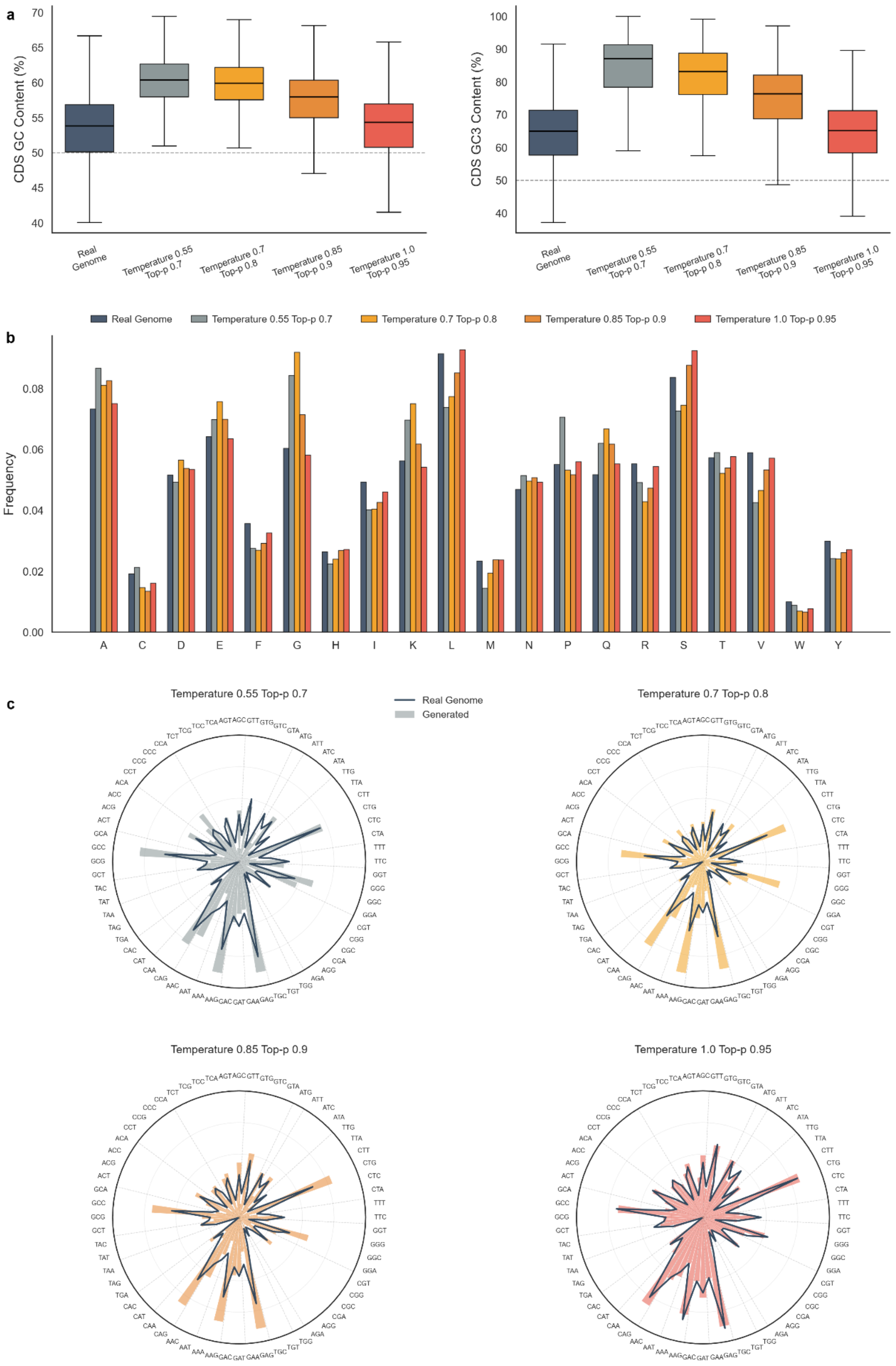

**Supplementary Fig. 7: Effects of sampling parameters on CDS features of generated *Drosophila melanogaster* sequences**

**a**, Distributions of CDS GC content (left) and GC3 content (right) for natural and generated sequences. **b**, Amino-acid usage preferences of natural and generated sequences. **c**, Codon usage preferences of natural and generated sequences across sampling conditions.
